# The Coordination and Jumps along C_4_ Photosynthesis Evolution in the Genus *Flaveria*

**DOI:** 10.1101/460287

**Authors:** Ming-Ju Amy Lyu, Udo Gowik, Peter Westhoff, Yimin Tao, Steve Kelly, Sarah Covshoff, Harmony Clayton, Julian M. Hibberd, Rowan F. Sage, Martha Ludwig, Gane Ka-Shu Wong, Xin-Guang Zhu

## Abstract

**Background:** C_4_ photosynthesis is a remarkable complex trait, elucidations of the evolutionary trajectory of C_4_ photosynthesis from its ancestral C_3_ pathway can help us to better understand the generic principles of complex trait evolution and guide engineering of C_3_ crops for higher yields. We used the genus *Flaveria* that contains C_3_, C_3_-C_4_, C_4_-like and C_4_ species as a system to study the evolution of C_4_ photosynthesis.

**Results:** We mapped transcript abundance, protein sequence, and morphological features to the phylogenetic tree of the genus *Flaveria*, and calculated the evolutionary correlation of different features. Besides, we predicted the relative changes of ancestral nodes of those features to illustrate the key stages during the evolution of C_4_ photosynthesis. Gene expression and protein sequence showed consistent modification pattern along the phylogenetic tree. High correlation coefficients ranging from 0.46 to 0.9 among gene expression, protein sequence and morphology were observed, and the greatest modification of those different features consistently occurred at the transition between C_3_-C_4_ species and C_4_-like species.

**Conclusions:** Our data shows highly coordinated changes in gene expression, protein sequence and morphological features. Besides, our results support an obviously evolutionary jump during the evolution of C_4_ metabolism.

## Background

Elucidating the evolutionary and developmental processes of complex traits formation is a major focus of current biological and medical research. Most health related issues, including obesity and diabetes, as well as agricultural challenges, such as flowering time control, crop yield improvements, and disease resistance, are related to complex traits [1-3]. Currently, genome-wide association studies are used to study complex traits. Putative genes or molecular markers of importance are then evaluated by a reverse genetics approach to identify those influencing the complex trait. C_4_ photosynthesis is a complex trait that evolved from C_3_ photosynthesis. When compared with C_3_ plants, C_4_ plants have higher water, nitrogen and light use efficiencies [4]. Interestingly, C_4_ photosynthesis has evolved independently more than 66 times, representing a remarkable example of convergent evolution [5]. Accordingly, C_4_ evolution is an ideal system for investigation of the mechanisms of convergent evolution of complex traits.

C_4_ photosynthesis contains a number of biochemical, cellular and anatomical modifications when compared with the ancestral C_3_ photosynthesis [6, 7]. In C_3_ photosynthesis, CO_2_ is fixed by ribulose-1,5-bisphosphate carboxylase/oxygenase (Rubisco), whereas in dual-cell C_4_ photosynthesis, CO_2_ is initially fixed into a four-carbon organic acid in mesophyll cells (MCs) by phospho*enol*pyruvate carboxylase (PEPC) [8]. The resulting four-carbon organic acid then diffuses into the bundle-sheath cells (BSCs) [9], where CO_2_ is released and fixed by Rubisco. Hence, C_4_ photosynthesis requires extra enzymes in CO_2_ fixation in addition to those already functioning in C_3_ photosynthesis, including PEPC, NADP-dependent malic enzyme (NADP-ME), and pyruvate, orthophosphate dikinase (PPDK) [8]. In dual-cell C_4_ photosynthesis, CO_2_ is concentrated in enlarged BSCs that are surrounded by MCs, forming the so-called Kranz anatomy [10-12]. Compared with C_3_ leaf anatomy, Kranz anatomy requires a spatial rearrangement of MCs and BSCs, cell size adjustment for increased numbers of organelles, larger organelles and metabolite transfer between the two cell types, and a reduction in distance between leaf veins.

Much of the current knowledge regarding the evolution of C_4_ photosynthesis was gained through comparative physiological and anatomical studies using genera that have not only C_3_ and C_4_ species, but also species performing intermediate types of photosynthesis [7, 13]. Among these, *Flaveria* has been promoted as a model for C_4_ evolution studies [14], and the evolution of C_4_-related morphological, anatomical and physiological features has been well studied in this genus over the last 40 years [14-17]. Though the molecular evolution of several key C_4_ enzymes were reported in this genus [18-20], however, the molecular evolution of C_4_ related features is large unknown. Besides, the evolutionary relationship of the C_4_ related molecular features and morphology features is not clear so far. In this study, we combined transcriptome data and published morphology data, together with the most recent phylogenetic tree of the genus *Flaveria* [21], to systematically investigate the key molecular events and evolutionary paths during the C_4_ evolution. Our results revealed that though many of the changes related to C_4_ photosynthesis occurred gradually, there are strong coordination and evolutionary jumps along the process.

## Results

### Transcriptome assembly and quantification

RNA-Seq data of 31 samples of 16 *Flaveria* species were obtained from the public database Sequence Read Achieve (SRA) of the National Center for Biotechnology Information (NCBI) (Table S1). The 16 species represented two C_3_ species, seven C_3_-C_4_ intermediate species, three C_4_-like species and four C_4_ species [22, 23] (Table S1). On average, 42,132 contigs (from 30,968 to 48,969) with lengths of no less than 300 bp were assembled for each of the 16 species. The N50 of these contigs ranged from 658 to 1208 (Table S2). The 16 species had a similar contig length distribution, with a peak at around 360 bp (Fig. S1). Since *Flaveria* is a eudicot genus, we used *Arabidopsis thaliana* (Arabidopsis) as reference to annotate *Flaveria* transcripts. On average, 58.91% of *Flaveria* contigs had orthologous genes in Arabidopsis.

Transcript abundance was calculated as fragments per kilobase of transcript per million mapped reads (FPKM) (see Methods). The total transcriptome-level comparison revealed higher Pearson correlations in overall transcript abundance in leaf samples from the same species than those of different organs from the same species, regardless of source (Fig. S2). Specifically, leaves from different developmental stages or from different labs are more closely correlated than leaf samples from different species, or than mean values of pair-wise correlations across all 27 leaf samples (T-test, P<0.05) (Fig. S3). Therefore, the mean FPKM value from multiple leaf samples was assigned as the final transcript abundance of the leaf for each species. Our quantification showed that, in *F. bidentis* (C_4_), transcripts from genes encoding C_4_-associated proteins were more abundant in leaf than in root and stem tissues (Fig. S4), which is consistent with earlier reports [24, 25]. In contrast, in the C_3_ species *F. robusta*, the difference in transcript abundance of orthologous genes between leaf and root/stem tissues was much less. Our data showed that NADP-ME is the dominant C_4_ pathway in *Flaveria* species (Fig. S5). We also proved that all species used in this study are from natural evolution and can be used for this evolutionary study (Fig. S6, see Supplementary Results). As a result, 13,081 Arabidopsis orthologues were detected in at least one of the 16 *Flaveria* species, and 12,215 genes were kept with the maximum FPKM in 16 species no less than 1 FPKM.

### C_4_ related genes: genes showed difference in gene expression and protein sequence between C_3_ species and C_4_ species

We first identified C_4_ related genes, which were defined as genes show difference in both gene expression and protein sequence between C_3_ and C_4_ species. We first calculated the differentially expressed (DE) genes between C_3_ and C_4_ species, which resulted in 2,306 DE genes. We next investigated transcriptome-wide amino acid changes predicted from orthologues of C_3_ and C_4_ *Flaveria* species using the process shown in Fig. S7 (Supplementary Methods). To estimate the accuracy of the predicted peptide sequences from our data, we conducted a comparative study of protein sequences from UniProtKB (http://www.uniprot.org) with those from our data, we found that our predicted peptide sequence is as good as, if not better than, the sequence from UniProtKB in terms of accuracy (details see Supplementary Results and Table S3). As a result, we obtained 1,018 genes encoding at least one amino acid change between C_3_ and C_4_ *Flaveria* species. 205 out of these 1,018 genes also showed significantly differentially expression (*P*<0.01) between C_3_ and C_4_ species, which was termed as C_4_ related genes, 113 and 92 showed ascending and descending transcript abundance in C_4_ species relative to C_3_ *Flaveria* species, respectively (Fig. S8).

We then investigated the degree of overlap of the 205 C_4_ genes from *Flaveria* with genes known or having the potential to be related to C_4_ photosynthesis or C_4_evolution in different species, including Arabidopsis [26], (Fig. S9), *Gynandropsis gynandra* [27] (Fig. S10), *Setaria viridis* [28, 29] (Fig. S11) and *Zea mays* (maize) [30] (Fig. S11 and Fig. S12). Result shows that the 205 genes are significantly enriched in those genes that are potentially related to C_4_ photosynthesis or C_4_ evolution (*P*<0.05, “BH” adjusted) (details see the Supplementary Results).

### The C_4_ related genes showed coordinated and abrupt change along the C_4_ evolutionary pathway in the genus *Flaveria*

The 205 genes are significantly enriched in several gene ontology (GO) terms including photosynthesis, photorespiration, photosynthetic light reaction, photosynthesis electron transport in photosystem I (PSI), photosynthesis light harvesting in PSI, chloroplast, organic anion transport and oxidation-reduction (*P*<0.05, *Fisher’s* exact test, “BH” adjustmented; Table S4). In the following sections, we systematically discuss these genes and their changes during C_4_ evolution in *Flaveria* with regard to gene expression and predicted protein sequences. We first focus on genes encoding proteins associated with C_4_ photosynthesis, then on genes related to the enriched GO terms (Table S4), which satisfy the following two criteria: (1) more than two predicted amino acid differences between C_3_ and C_4_ species, and (2) fully assembled predicted protein sequences from species belonging to all four photosynthetic types: C_3_, C_3_-C_4_, C_4_-like and C_4_. In general, these genes were classified into six categories according to the probable function of their cognate proteins, *i.e*., the C_4_ pathway, photorespiratory pathway, electron transport chain, membrane transport, photosynthetic membrane, and oxidation-reduction.

#### Genes encoding proteins associated with the C_4_ pathway

Nine genes encoding proteins associated with the C_4_ pathway were identified, including those encoding three C_4_ cycle enzymes, PEPC, PPDK and NADP-ME, two regulatory proteins, PPDK regulatory protein (PPDK-RP) and PEPC protein kinase A (PPCKA), two aminotransferases, Alanine aminotransferase (AlaAT) and aspartate aminotransferase 5 (AspAT5), and two transporters, BASS2 and sodium: hydrogen (Na^+^/H^+^) antiporter 1 (NHD1) (Table 1). In terms of protein sequence, the major predicted amino acid changes in C_4_ species occurred at N7 for all of the nine genes (Figs. 1, 2, Figs. S13, Table 1). For example, PEPC in the C_4_ *Flaveria* species had 41 predicted amino acid changes compared with those in the C_3_ species, which were mapped onto the *Flaveria* phylogeny determined by Lyu *et al*. [21]. One of the predicted changes occurred at N6 (D396 in C_4_ species, hereafter D396), and 34 occurred at N7 (Fig. 1A). The six other predicted amino acid changes occurred at N7 or after N7, although the incomplete assembly of PEPC transcripts from *F. palmeri* and *F. vaginata* did not allow resolution of the predicted amino acid sequences. These results suggest an evolutionary jump in the protein sequence at N7 for C_4_ enzymes.

**Figure 1.**
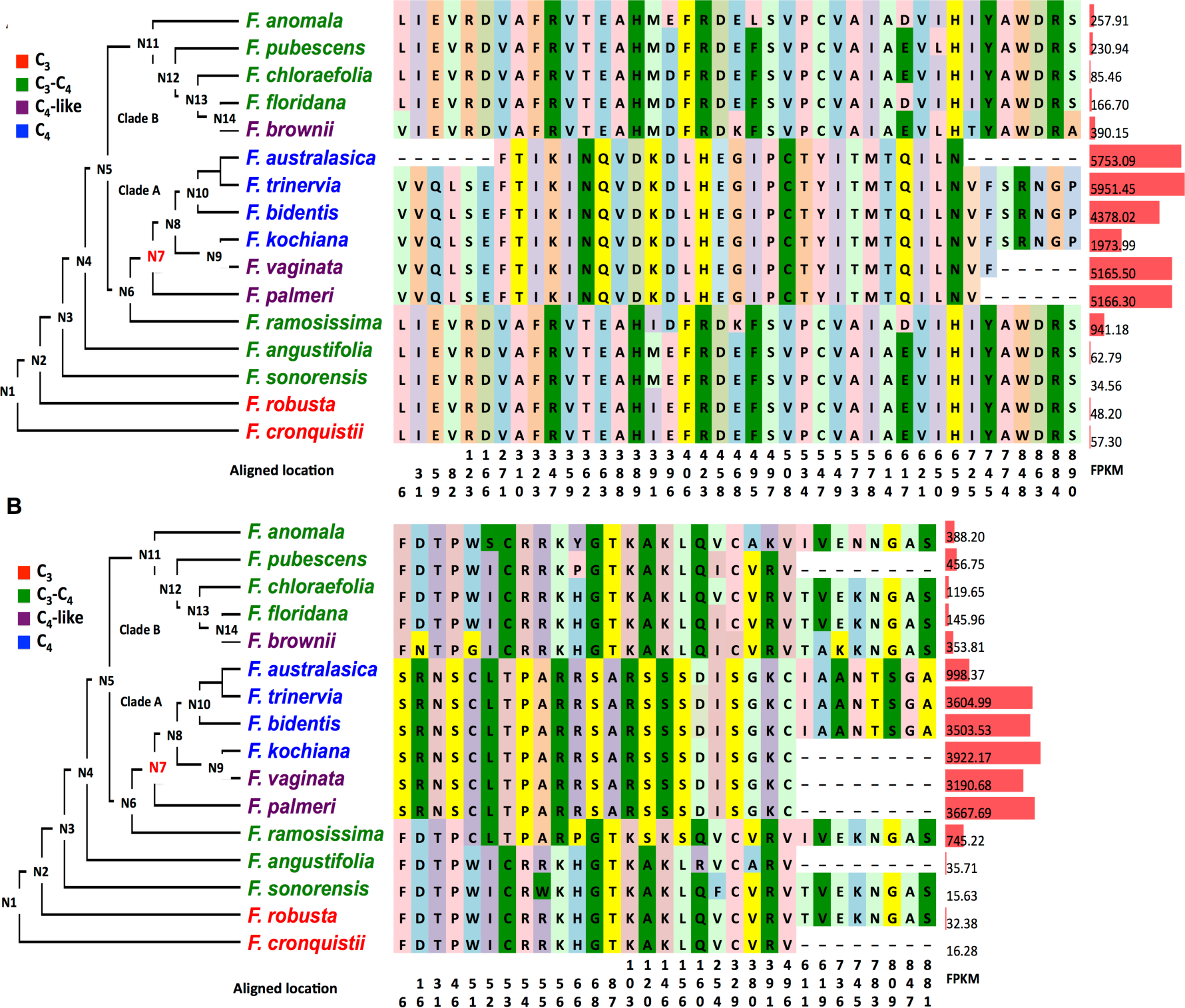
Modifications in phospho*enol*pyruvate carboxylase and pyruvate orthophosphate dikinase predicted protein sequences and transcript abundances mapped to the *Flaveria* phylogeny. The predicted amino acid changes in phospho*enol*pyruvate carboxylase (PEPC) and pyruvate orthophosphate dikinase (PPDK) between C_4_ and C_3_ *Flaveria* species and the transcript abundance (FPKM) of the genes encoding the proteins are shown. Only the amino acid residues predicted to be different between C_3_ and C_4_ species are superimposed on the schema of *Flaveria* phylogenetic tree modified from Lyu *et al.*, 2015. The colors of amino acid residues have no meaning and are only for visualization purposes. Numbers below the amino acids indicate the location sites in the multiple sequence alignments. FPKM values are shown to the right of the amino acid changes as red bars. A: PEPC. B: PPDK. Protein sequences from UniprotKP are: *F. trinervia* PEPC, P30694; *F. bidentis* PPDK, Q39735; *F. brownii* PPDK, Q39734; and *F. trinervia* PPDK, P22221. Sequence alignments are available in Additional file 2.

**Figure 2.**
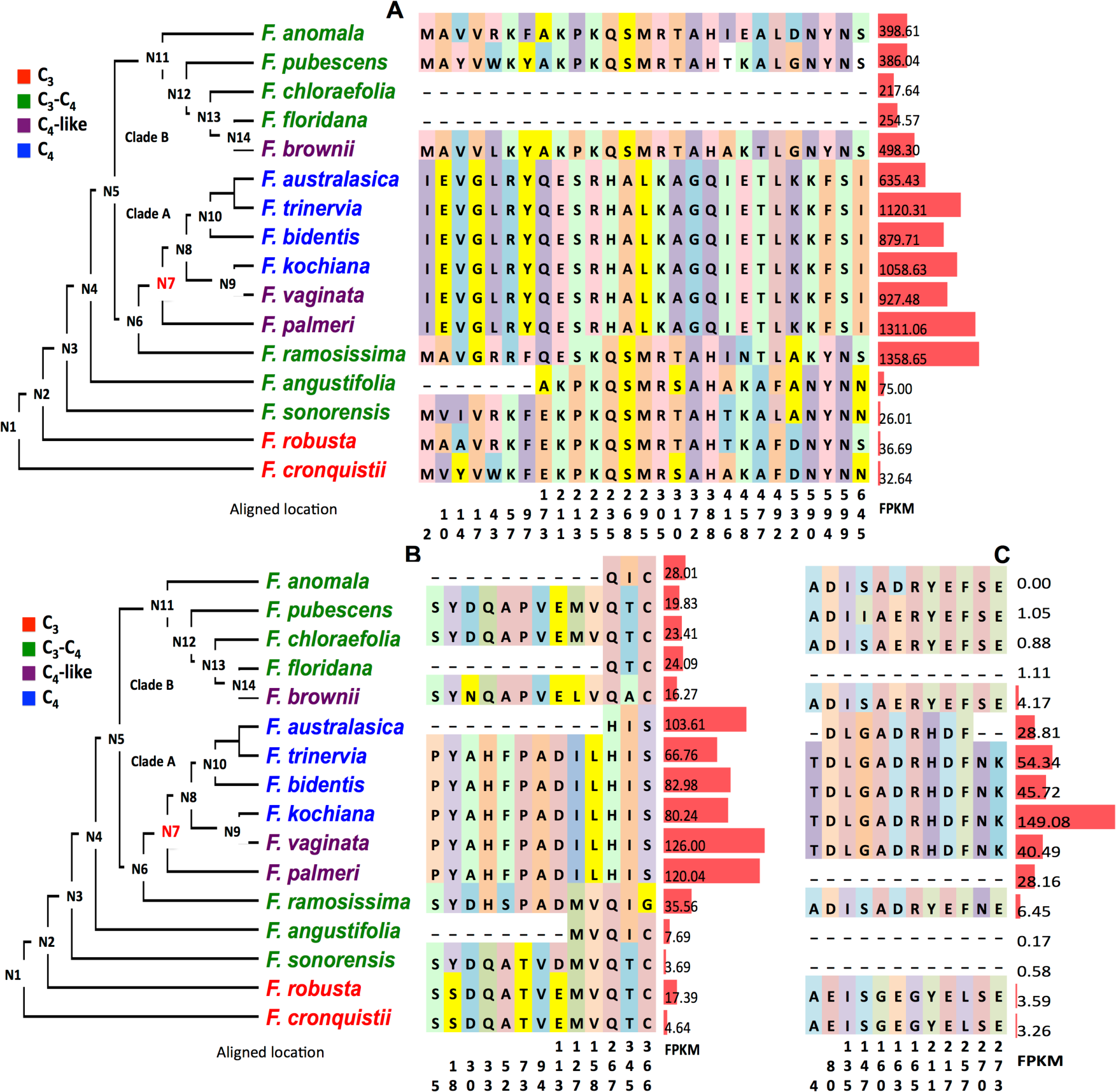
Modifications in NADP-malic enzyme, pyruvate orthophosphate dikinase regulatory protein and phospho *enol* pyruvate protein kinase A predicted protein sequences and transcript abundances mapped to the *Flaveria* phylogeny. The predicted amino acid changes in NADP-malic enzyme (NADP-ME), pyruvate orthophosphate dikinase regulatory protein (PPDK-RP) and phospho*enol*pyruvate protein kinase A (PPCKA) between C_4_ and C_3_ species and the transcript abundance (FPKM) of the genes encoding the proteins are shown. Only the amino acid residues predicted to be different between C_3_ and C_4_ species are superimposed on the schema of *Flaveria* phylogenetic tree modified from Lyu *et al.*, 2015. The colors of amino acid residues have no meaning and are only for visualization purposes. Numbers below the amino acids indicate the location site in the multiple sequence alignments. FPKM values are represented to the right of the amino acid changes as red bars. A: NADP-ME. B: PPDK-RP. C: PPCKA. The sequence alignments are available in Additional file 2.

**Table 1.**
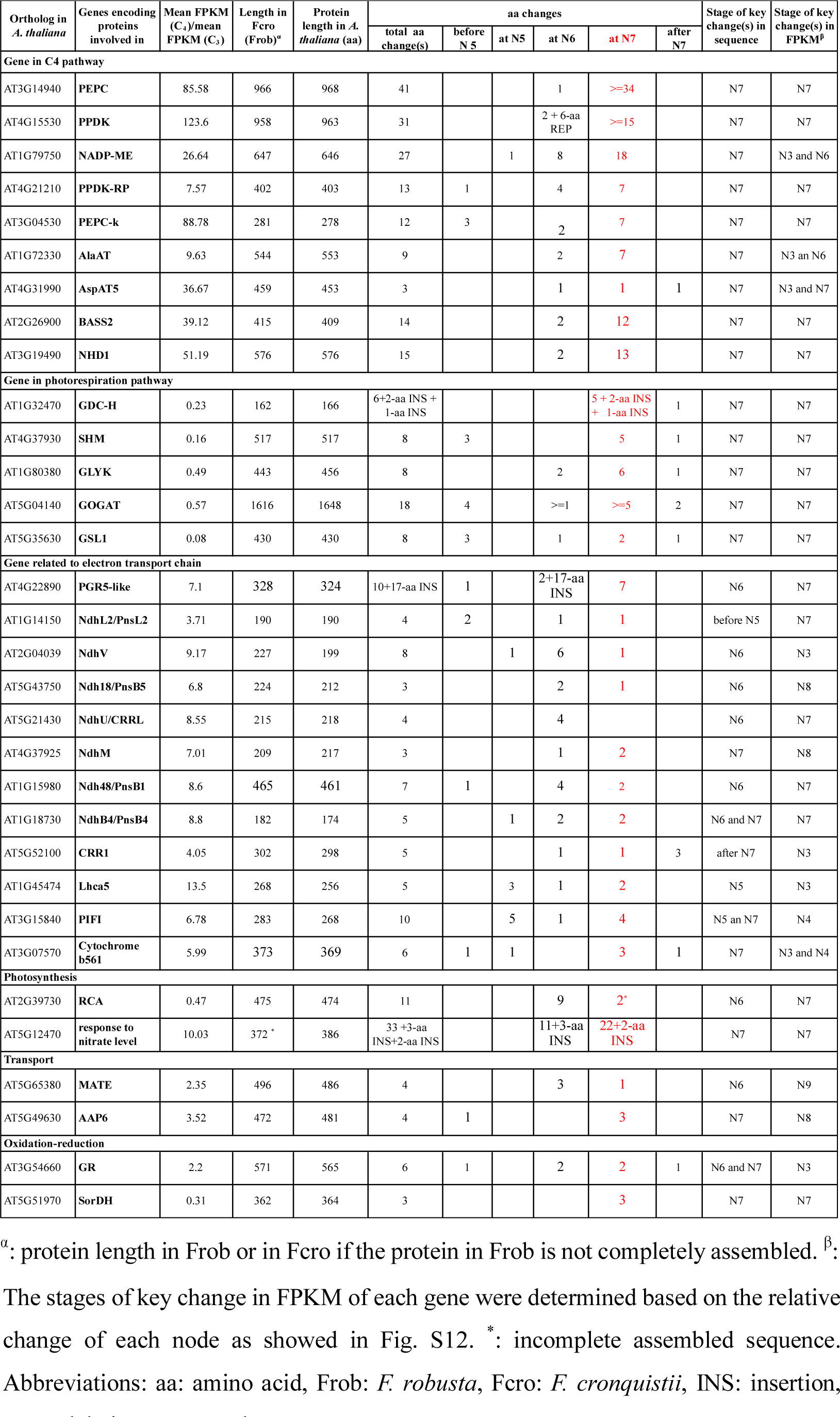
Proteins showing differences in amino acid sequence between C_3_ and C_4_ *Flaveria* species and the relative changes in their cognate transcripts

In terms of gene expression, all the nine genes showed higher transcript abundance in C_4_ species than in C_3_ species and a comparable level in C_4_-like and C_4_ species (Table 1). To calculate the relative gene expression changes of each ancestral node, the FPKM values of each ancestral node were predicted and the relative difference were calculated (see Methods). In general, C_4_ species showed a 7.6-fold to 123.6-fold of FPKM values compared with C_3_ species. Similar to the pattern of changes for protein sequences, seven of the nine genes showed that the biggest relative changes of gene expression at N7. Whereas, both NADP-ME and AlaAT showed the biggest relative changes at two nodes of N3 and N6 with comparable levels (Fig. 2A and Fig. S13A). Our results hence suggested that the genes encoding proteins associated with C_4_ pathway showed highly coordinated modification patterns in protein sequence and gene expression at N3, N6 and N7 during the evolutionary pathway of C_4_ photosynthesis; while the majority of the predicted amino acid changes occurs at the N7.

#### Genes encoding proteins in the photorespiratory pathway

Transcripts encoding five proteins involved in photorespiration were identified in our comparative analyses: glycine decarboxylase complex (GDC) H subunit (GDC-H), serine hydroxymethyltransferase (SHM), glycerate kinase (GLYK), glutamine synthetase and glutamine oxoglutarate aminotransferase (GOGAT), and glutamine synthetase-like 1 (GSL1) (Figs. 3, Table 1). In general, the predicted amino acid substitution patterns of these five proteins were similar to those observed in the above described proteins in C_4_ pathways, with the major predicted amino acid changes in C_4_ species occurring at N7 (Figs 3, Table 1), *e.g.,* 16 of 18 in GOGAT occurred at N7 (Fig 3D). Generally, proteins in the photorespiratory pathway showed fewer predicted amino acid changes than those in the C_4_ pathway.

**Figure 3.**
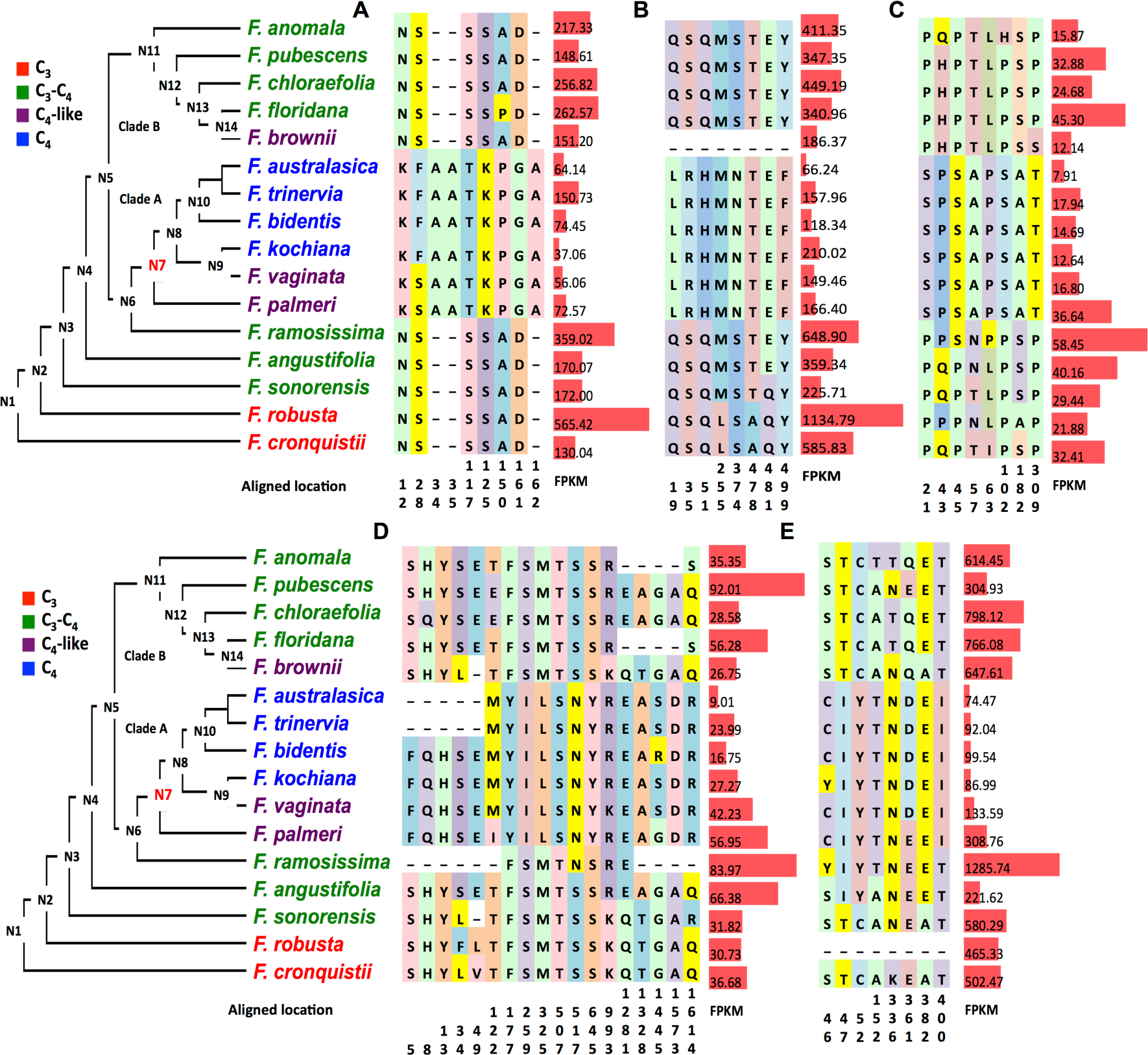
Modifications in photorespiratory protein predicted amino acid sequences and cognate transcript abundances mapped to the *Flaveria* phylogeny. The predicted amino acid changes in photorespiratory proteins between C_4_ and C_3_ *Flaveria* species and the transcript abundance (FPKM) of genes encoding the proteins are shown. Only the amino acid residues that are predicted to be different between C_3_ and C_4_ species are superimposed on the schema of *Flaveria* phylogenetic tree modified from Lyu *et al.*, 2015. The marked colors of amino acid residues have no meaning and are only for visualization purposes. Numbers below the amino acids indicate the location site in the multiple sequence alignments. FPKM values are represented to the right of the amino acid changes as red bars. A: glycine decarboxylase complex H subunit (GDC-H); B: serine hydroxymethyltransferase (SHM); C: glycerate kinase (GLYK); D: glutamine synthetase and glutamine oxoglutarate aminotransferase (GOGAT); E, Glutamine synthetase like 1 (GSL1).

The abundance of transcripts encoding the five photorespiratory enzymes examined was comparable in C_3_ and C_3_-C_4_ species, and higher than in the C_4_ species (Figs. 3 A–E). Interestingly, we found a consistent pattern in transcript abundance that paralleled the trajectory of C_4_ evolution, with the highest, or the second highest FPKMs found in the C_3_-C_4_ species *F. ramosissima* and the lowest observed in the C_4_ species. For example, the FPKM values for GOGAT were 30.73 in *F. robusta* (C_3_), 83.97 in *F. ramosissima* (C_3_-C_4_), 56.95 in *F. palmeri* (C_4_-like) and 27.27 in the C_4_ species *F. kochiana* in clade A. In clade B, the values changed from 30.73 in the C_3_ *F. robusta* to 35.35 in *F. anomala* (C_3_-C_4_), 92.01 in *F. pubescens* (C_3_-C_4_), and 26.27 in *F. brownii* (C_4_-like). Moreover, transcripts encoding photorespiratory enzymes exhibited at least a 1.5-fold difference between C_4_-like species and C_4_ species in clade A. Specifically, when compared with *F. kochiana* (C_4_), *F. vaginata* (C_4_-like) showed a 1.55-fold increase for GOGAT (27.27 to 42.23), a 1.54-fold increase for GSL1 (86.99 to 133.59), and a 1.51-fold increase for GDC-H (37.06 to 56.06). *F. palmeri* displayed an even larger fold-change relative to *F. kochiana* than *F. vaginata* (Figs. 3 A-E). Hence, when compared with genes encoding C_4_ pathway proteins, those encoding photorespiratory proteins showed larger differences between C_4_-like and C_4_ species in clade A in terms of gene transcript abundance and cognate protein sequence. The greatest reduction of FPKM was observed uniformly at N7 (Figs. 3, Table 1). Thus, this suggested that the genes encoded proteins associated with photorespiratory pathway also showed coordinated changing pattern in protein sequence and gene expression during the evolutionary pathway of C_4_ photosynthesis, again with the largest number of changes occurring at N7.

#### Genes encoding proteins involved in the electron transport chain

We identified genes encoding 12 proteins that function in the photosynthetic electron transfer chain, including nine related to cyclic electron transport (CET) (Fig. 4), PSI light harvesting complex gene 5 (Lhca5), post-illumina chlorophyll fluorescence increase (PIFI) and cytochrome b561 (Cytb561). Transcripts encoding all 12 of the proteins showed higher abundances in C_4_ species than C_3_ species (Figs. 4, Table 1). The genes encoding proteins involved in CET showed the biggest changes at different nodes instead of at a single node, *e.g.,* the major changes of PGR5-like in FPKM and in predicted protein sequences occurred at N6 and N7, respectively (Fig. 4A). Transcripts encoding NdhL2 showed the biggest increase in abundance at N7, and two of four predicted amino acid changes occurred before N5 (Fig. 4B); the major changes of Lhca5 were observed at N5 (Fig. S14A). This suggested that the variation of CET might have contributed to split of clade A and B in the *Flaveria* genus.

**Figure 4.**
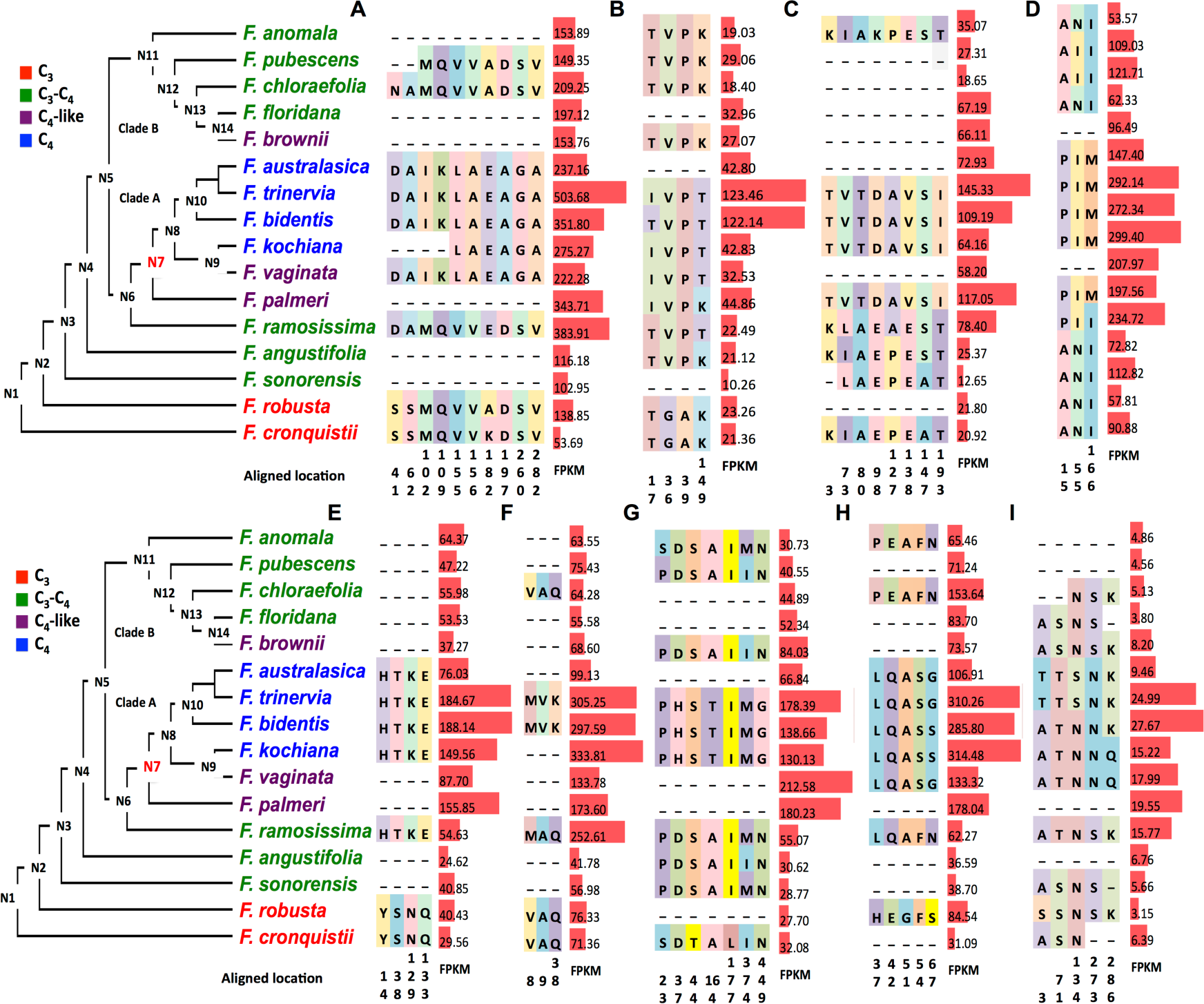
Modifications in the predicted amino acid sequences of proteins involved in cyclic electron transport and transcript abundances of the cognate transcripts mapped to the *Flaveria* phylogeny. Changes in predicted amino acid sequence in proteins involved in cyclic electron transport and abundances (FPKM) of their cognate transcripts in C_4_ and C_3_ *Flaveria* species are shown. Only the amino acid residues predicted to be different between C_3_ and C_4_ species are superimposed on the schema of *Flaveria* phylogenetic tree modified from Lyu *et al.*, 2015. The marked colors of amino acid residues have no meaning and are only for visualization purposes. Numbers below the amino acids indicate the location site in the multiple sequence alignments. FPKM values are represented to the right of the amino acid changes as red bars. A: protein gradient regulation 5 like protein (PGR5-like); B: NADH dehydrogenase-like (Ndh) L2 subunit (Ndh L2); C: NdhV; D: Ndh16; E: NdhU; F: NdhM; G: Ndh48; H: NdhB4; I: chlororespiration reduction 1. The sequence alignments are available in Additional file 2.

#### Genes encoding proteins associated with photosynthesis, transport, and oxidation-reduction

Two genes labelled in GO term as photosynthesis, namely, Rubisco activase (RAC) and orthologue of AT5G12470 were identified. RAC showed the greatest decrease in FPKM at N7, while nine out of eleven predicted modifications in its sequence were acquired at N6 (Fig. S15A, Table 1). The orthologue of AT5G12470 in *Flaveria*, (hereafter *Flaveria*-AT5G12470), a chloroplast envelope protein [31] potentially involved in responses to nitrate levels [32], showed the greatest changes in transcript abundance and predicted protein sequences at N7 (Fig. S15B, Table 1).

Transcripts encoding two transport proteins also displayed changes in FPKM and predicted amino acid sequence in C_4_ species and C_3_ *Flaveria* species, namely, multidrug and toxic compound extrusion (MATE) protein and amino acid permease 6 (AAP6) (Fig. S16A and B). AAP6 was reported to be a high affinity neutral amino acid transporter primarily expressed in sink tissue and xylem parenchyma cells, and potentially responsible for taking up amino acids from the xylem and delivering them to the phloem [33-35]. The modifications in expression levels were observed at N9 for MATE and N8 for AAP6, and the greatest modifications in predicted sequence were observed at N6 and N7, respectively (Fig. S16, Table 1).

Transcripts encoding two proteins playing roles in oxidation-reduction showed changes between C_4_ and C_3_ species of *Flaveria*, namely, glutathione reductase (GR) and sorbitol dehydrogenase (SorDH). GR showed the major enhancement of transcript abundance at N3 and predicted amino acid changes at N6 and N7 (Fig. S17A, Table 1), Similarly, the major reduction of transcript abundance of and predicted amino acid changes of SorDH showed at N7 (Fig. S17B). These results are consistent with studies suggesting a pivotal role of redox status in the expression of genes encoding photosynthetic proteins [36, 37].

### Physiological and anatomical characteristics related to C_4_ photosynthesis show coordinated changes along the C_4_ evolutionary pathway in *Flaveria*

To investigate whether C_4_-related physiological characteristics also underwent coordinated changes during the evolution of C_4_ photosynthesis in *Flaveria*, physiological characteristics taken from the literature [23, 38, 39] were mapped onto the *Flaveria* phylogeny (Fig. 5). The results revealed a step-wise change for most of the characteristics along the phylogenetic tree as previously suggested [14, 23, 38, 39] (Fig. 5); however, coordinated and abrupt changes were observed for a number of features. A major change in CO_2_ compensation point (Γ) in *Flaveria* was first seen at N3, where the most ancestral C_3_-C_4_ species, *F. sonorensis*, was emerged which showed a decrease in Γ from 62.1 μbar of its closest C_3_ relative *F. robusta* to 29.6 μbar (Fig. 5). The greatest changes in Γ in clade A occurred at N6, which showed a decrease in Γ from 24.1 μbar in *F. angustifolia* (C_3_-C_4_) to 9.0 μbar in *F. ramosissima* (C_3_-C_4_), followed by N7, where a decrease in Γ from 9.0 μbar in *F. ramosissima* (C_3_-C_4_) to 4.7 μbar in *F. palmeri* (C_4_-like) was seen. The greatest decrease of Γ in clade B was observed between the two C_3_-C_4_ species, *F. floridana* and *F. chloraefolia* (C_3_-C_4_), where there was a decrease from 29 μbar to 9.5 μbar (Fig. 5). For photosynthetic water using efficiency (PWUE), photosynthetic nitrogen using efficiency (PNUE) and the slope of the response of the net CO_2_ assimilation rate (*A*) versus Rubisco, the biggest changes occurred at N7 with increases of around 2-fold. In contrast, the percentage of ^14^C fixed into four carbon acids showed no clear trend along the phylogenetic tree, although 3.91-fold and 1.76-fold increases were seen at N6 and N7, respectively. Interestingly, changes in all of these traits uniformly occurred at *F. brownii* in clade B, the only C_4_-like species within this clade. Consequently, those data suggest that although there were gradual changes in physiological features along the C_3_, C_3_-C_4_, C_4_-like and C_4_ trajectory, there are apparent jumps at N3, N6 and N7 in these physiological traits along the *Flaveria* phylogeny (Fig. 5).

**Figure 5.**
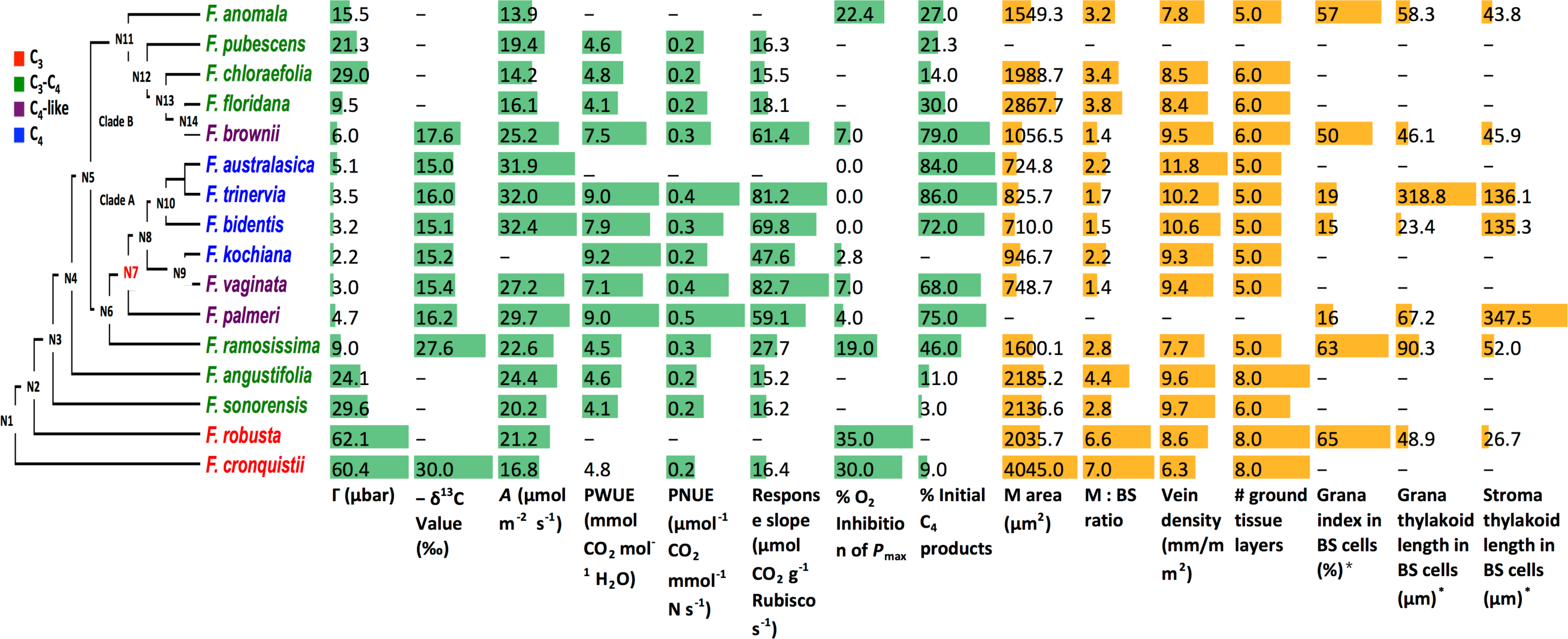
Changes in physiological and anatomical traits mapped onto the *Flaveria* phylogeny. Overall, C_4_-related physiological (green and blue bars) and anatomical traits (orange and red bars) showed a step-wise change along the *Flaveria* phylogenetic tree; however, a number of the traits showed greater more significant changes at certain nodes. *Grana index: total length of grana/total length of thylakoid membrane X 100. (Abbreviations: Γ: CO_2_ compensation point; *A*: CO_2_ assimilation rate; PWUE: instantaneous photosynthetic water use efficiency; PNUE: instantaneous photosynthetic nitrogen use efficiency; response slope: slope of the response of net CO_2_ assimilation rate versus leaf Rubisco content; M: mesophyll; BS: bundle sheath.) Data are from references as given in the Methods.

Anatomical traits [14] were mapped onto the *Flaveria* phylogeny to investigate how these features were modified along the evolution of C_4_ (Fig. 5). For both the area of MCs and the ratio of the area of MCs to that of BSCs (M: BS), the greatest modifications along the phylogeny were found between *F. brownii* (C_4_-like) and *F. floridana* (C_3_-C_4_), with a similar degree of change for both characteristics (2.7-fold, Fig. 5). Anatomical data for *F. palmeri* (C_4_-like) in clade A are not available; however, large differences in anatomical features were found between the C_4_-like *F. vaginata* and C_3_-C_4_ *F. ramosissima* [14]. The modification of M area first occurred at N2 which showed a 1.9-fold difference (compare 4045 m^2^ in *F. robusta* to 2035.7 m^2^ in *F. cronquistii*) followed by a 2.1-fold of difference between *F. ramosissima* and *F. vaginata* (compare 1600.1 m^2^ in *F. ramosissima* and 748.7 m^2^ in *F. vaginata*). The major modification of the ratio of M and BS occurred at N3 with a 2.4-fold difference (compare 6.6 m^2^ in *F. robusta* and 2.8 m^2^ in *F. sonorensis*) and N6 with a 1.6-fold difference (compare 4.4 m^2^ in *F. angustifolia* and 2.8 m^2^ in *F. ramosissima*) and between *F. ramosissima* and *F. vaginata* (1.4 m^2^) with a 2-fold difference. Therefore, similar to the evolutionary pattern of physiological features, large changes in anatomical features also emerged at N3, N6 and the transition between C_3_-C_4_ species and C_4_-like species. Interestingly, the ultrastructure of BSCs chloroplasts showed an abrupt change at N7, with a dramatic decrease in grana thylakoid length and an increase in stroma thylakoid length, whereas these features were comparable in the species at the base of tree and in clade B [40]. These findings are consistent with the observation that the abundance of transcripts encoding proteins involved in CET increased more in clade A species than in clade B species, as described above. These results imply that CET only increased in clade A species, which may be a key factor determining the possibility of forming C_4_ species from the C_3_-C_4_ intermediate species.

### Coordinated change of protein sequence, gene expression and morphology with an evolutionary jump at the transition between C_3_-C_4_ and C_4_-like species along the species evolution

Our above analysis showed that C_4_ related features showed coordinated changes with an obvious abrupt change at N7. Next, we asked whether species evolution also showed evolutionary coordination and jump(s) along the species evolution in protein sequence, gene expression and morphology. To answer this question, we calculated the divergence matrics for protein sequence, gene expression, and morphological features between *F. cronquistii* (at the most basal place in the *Flaveria* phylogenetic tree) and other *Flaveria* species. The protein divergence was calculated as the rate of non-synonymous substitutions (dN) of all the genes that were used to construct the *Flaveria* phylogenetic tree from [21]; and expression divergence as Euclidean distance of total expressed genes (see Methods); and morphology divergence as Euclidean distance using previously coded morphology value from [41], which includes 30 types of morphology traits, such as life history, leaf shape, head types and so on. Our data showed a high linear correlation between the protein divergence, gene expression divergence and morphology divergence, in particular between gene expression divergence and morphology divergence (R^2^=0.9) (Figs. 6A-C), suggesting a coordinated evolution of protein sequence, gene expression and morphology in species evolution.

**Figure 6.**
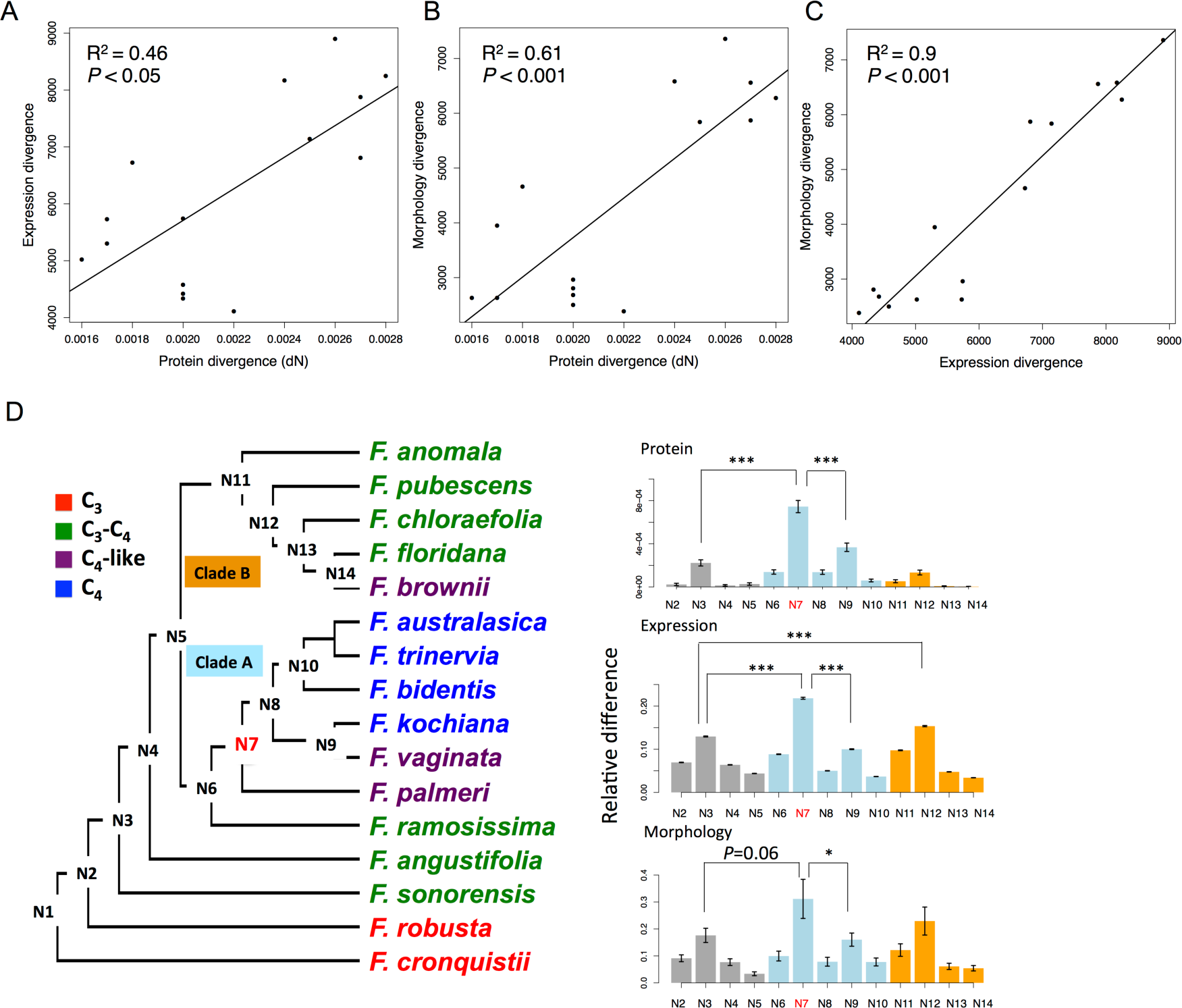
Coordinated evolution of protein sequence, gene expression and morphology with an obvious jump change. Significant linear correlation between protein divergence, gene expression divergence and morphology divergence were showed in (A-C). Protein divergence was calculated as non-synonymous mutation (dN). Expression divergence and morphology divergence were calculated as Euclidean distance based on quantile normalized FPKM values and coded morphology values from Mckown *et al.*, 2005, respectively. All the Mckown relative divergences were the divergence between *F. cronquistii* and other *Flaveria* species. (D) Shows the relative difference of each ancestral node compared with its earlier ancestral node in protein sequence, gene expression and morphology. The left panel shows the schema of *Flaveria* phylogenetic tree modified from Lyu, *et al*., 2015. Each ancestral node was numbered according to the evolutionary time. *P* values are from One-way ANOVA analysis followed by Tukey’s Post Hoc test and adjusted by *Benjamin-Hochberg* correction. The significant levels are: *: *P*<0.05; **: *P*<0.01; ***: *P*<0.001. The bar colors in grey/blue/orange represent species from basal/clade A/clade B of phylogenetic tree, respectively.

Next, we predicted the protein sequence, transcript abundance and coded morphology value of ancestral nodes, which were then used to calculate the relative change of the three parameters at each node (see Methods). Surprisingly, protein sequence and gene expression showed significantly more changes at N7 than changes at other nodes (*P*<0.001, Tukey’s test, “BH” adjusted , the same as following), and the morphology showed the most changes at N7 with a marginal significant *P* value (*P*=0.06), implying a evolutionary coordination and jump also occurred in species evolution.

Consider that N7 is the most recent common ancestor of C_4_ and C_4_-like species in clade A, where C_4_-like and C_4_ species are comparable with respect to the C_4_-ness (Figs. 6), it may be possible that the C_4_ photosynthesis accelerate the evolution of species. We then investigate how much total species variances can be explained by C_4_ related genes. Principle component analysis (PCA) showed that species derived from N7 were distinguished from other species (Figs. 7), which is consistent with the evolutionary jump at N7. The first component of the 205 C_4_ related genes account for 38% of total variance (Fig. 7C), more than the dataset of genes that expressed in all species (8004 genes), which account for 32% of total variance (Fig. 7A), and the DE genes account for 27% of total variance (Fig. 7B). Moreover, the 205 C_4_ related genes showed same evolutionary pattern with the total expressed genes and DE genes, which had the biggest changes at N7 (Figs. 7) (*P*<0.001), This raises the possibility that the evolution of species in the *Flaveria* genus might be mainly driven by the evolution of C_4_ photosynthesis. This is not surprising considering the generally higher light, nitrogen and water use efficiency in C_4_ photosynthesis as compared with C_3_ photosynthesis. It is highly likely that many parameters related to growth, development and responses to environments differ between species with these two different photosynthetic pathways.

**Figure 7.**
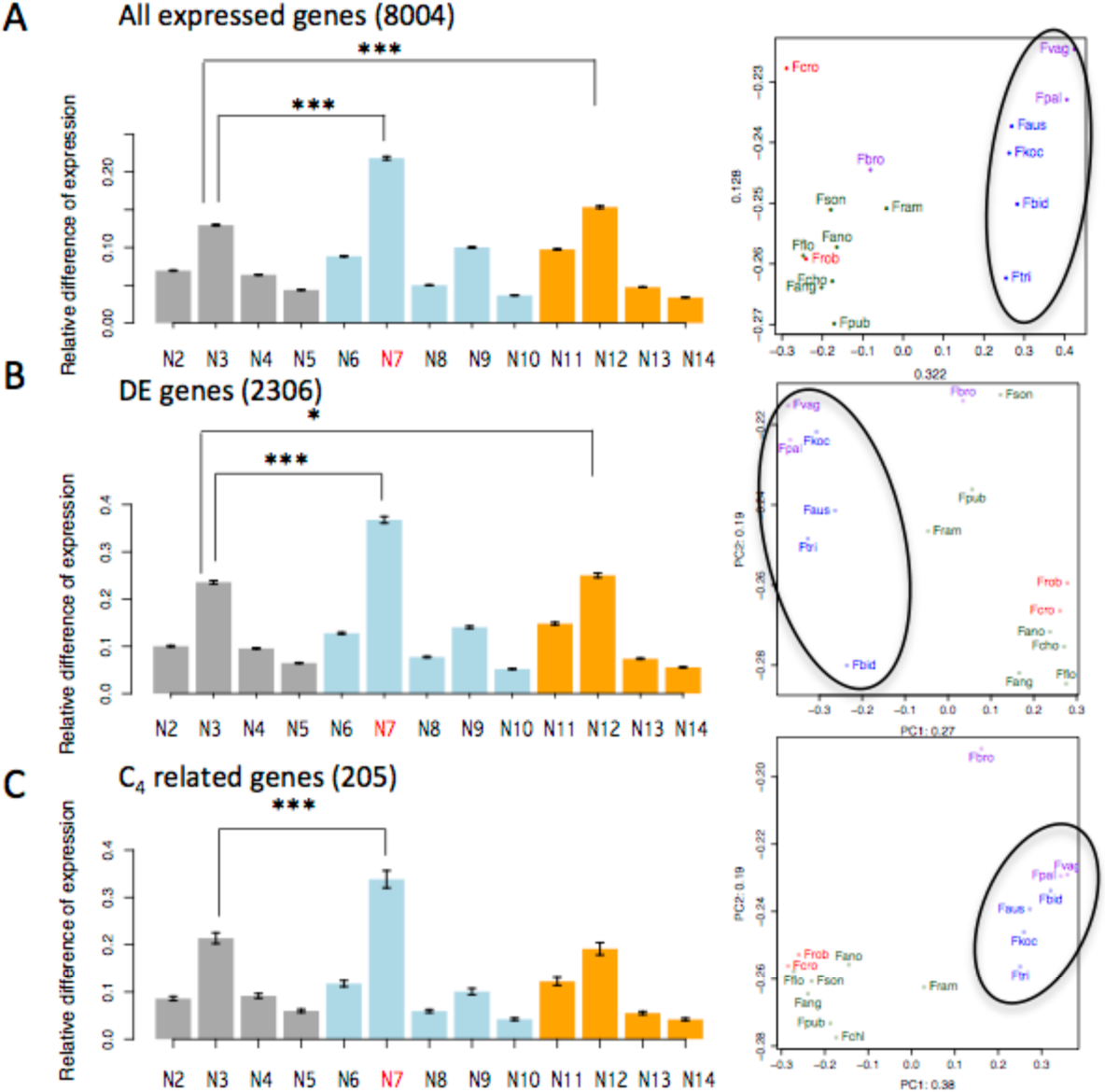
Evolutionary pattern of C_4_ related genes comparing with total expressed genes and DE genes between C_3_ and C_4_ species in gene expression. There dataset are (A) the total expressed gene (12215 genes), (B) differentially expressed genes between C_3_ and C_4_ species (2306 genes) and (C) C_4_ related genes that showed difference between C_3_ and C_4_ species in both protein sequence and gene expression (205 genes). All the three datasets showed that the biggest change of gene expression occurred at N7 (left panels). Principle Component Analysis (PCA) showed that species derived from N7 (species in the black frames in right panels) are distinguished with other species, and the first component of the C_4_ related genes account for 38% of the total variance, more than that of total expressed genes and DE genes (right panels). The bar colors grey/blue/orange represent: species from basal/clade A/clade B of phylogenetic tree.

## Discussion

### Evolutionary coordination of different features implies a purifying selection towards a functional C_4_ metabolism

Compared with C_3_ photosynthesis, C_4_ photosynthesis acquired many new features in gene expression, protein sequence, morphology and physiology (Figs. 1-6) [42]. We interpret these coordinated changes as a result of a strong purifying selection at this step. This is because though C_4_ photosynthesis can gain higher photosynthetic energy conversion efficiency, highly specialized leaf and cellular anatomical features and biochemical properties of the involved enzymes are required. For example, increased cell wall thickness at the bundle sheath cell and decreased sensitivity of PEPC to malate inhibition are needed for C_4_ plants to gain higher photosynthetic rates [43, 44]. Furthermore, to gain higher photosynthetic efficiency in C_4_ plants, the ratio of the quantities of Rubisco content in BSCs and MCs is also critical [45]. In theory, if the C_4_ decarboxlyation is in place occurs before all other accompanying features, leaves will experience high leakage, *i.e.*, costing ATP for a futile cycle without benefit to CO_2_ fixation. This will inevitably lead to lower quantum yield and a potential driving force for purifying selection. Further evidence for possible purifying selection comes from the observation that genes with cell-specific expression, such as PEPC, PPDK, and NADP-ME, displayed more changes in their predicted protein sequences than ubiquitously expressed genes, such as NDH components (Table 1, Additional file 3). This is because, as discussed earlier, the redox environments between BSCs and MCs might have changed dramatically during the completion of the C_4_ cycle, with one of the most likely change being having a more acidic environment due to increased production of Oxaloacetic acid (OAA) and malate. Under such conditions, it is required for enzymes to alter their amino acid sequences to adapt to the new cellular environments. The concurrent changes between gene expression divergence and protein sequence divergence has also been demonstrated previously in animals [46, 47], which has been similarly proposed to reflect negative selection for the involved genes [47].

### Evolutionary jumps along the C_4_ evolution in the *Flaveria* genus

Among the nodes leading to the C_4_ emergence in clade A, the N7 shows the biggest change in protein sequence, gene expression and morphology in both C_4_ specific features and also general features (Table 1, Figs. 1-7). There are also apparent changes in these features at N3 and N6. These three nodes reflect three critical stages during the emergence of C_4_ metabolism. First, at N3, there was a large degree of changes in gene expression, protein sequence and morphology (Figs. 1 and 2). One of the most important events during this phase is the re-location of GDC from MSCs to BSCs based on earlier western blot data [13, 48]. Here we found that SHM showed decreased expression while most of other photorespiratory related enzymes showed little changes (Figs. 4). Similarly, at this step, the majority of the C_4_ related genes showed little changes (Figs. 1 and 2). In contrast, global survey of the gene expression, protein sequence and morphology changes suggest that there is large number of changes at this step compared to earlier C_3_ species (Figs. 6 and 7), and there is also great decrease of CO_2_ compensation point at this stage (Fig. 5).

N6 is the stage where we found the third largest degree of changes occurred in C_4_ related features. At this stage, we observed large increase in transcripts coding for nearly all enzymes involved in nitrogen rebalancing (Figs. 1-3), photorespiration related transcripts, and concurrent increase in transcripts encoding the other remaining C_4_ cycle-related enzymes, and a dramatic increase in the percentage of ^14^C incorporated into the four carbon acids occurred (Fig. 5). The increase in transcript abundance in photorespiratory genes might be related to the optimization of C_2_ cycle to decrease CO_2_ concentrating point, which can increase fitness of plants under conditions favoring photorespiration [49]. The dramatic increase in enzymes related to nitrogen rebalancing, i.e. PEPC, NADP-ME, PEPCKA etc, is consistent with the notation that C_4_ cycle might be evolved as a result of rebalancing nitrogen metabolism after GDC moving from MC to BSC [17]. The fact that there is little change in the δ^13^C in the C_3_-C_4_ intermediate as compared to that of C_3_ species suggests that the contribution of CO_2_ fixation following C_4_ pathway is relatively minor, *i.e.*, less than 15% estimated based on an δ^13^C value of −27.6 in *F. ramosissima* (Fig. 5), again supporting the initial role of increased C_4_ enzymes is not for enhancing CO_2_ fixation. It is worth pointing out here that the measured initial carbon fixation in the form of C_4_ compound was 46% (Fig. 5), higher than those estimated based on the δ^13^C value. This is possibly because though malate releases CO_2_ into BSCs as a result of the nitrogen rebalancing pathway, most of the CO_2_ was not fixed by Rubisco, either due to lack of sufficient Rubisco activity in BSCs or due to lack of required low BSCs cell wall permeability to maintaining high CO_2_ concentration in BSCs *etc*.

N7 witnesses abrupt changes for both the gene expression and proteins sequence and morphology (Figs. 1-5, Fig. 6 D). The majority of the C_4_ related genes showed the most modification in gene expression and protein sequence at N7, especially for genes in C_4_ cycle and photorespiratory pathway. Moreover, N7, where C_4_-like species (clade A) appear, represents a dramatic shift of CO_2_ fixation from being dominated by a C_2_ concentrating mechanism to being dominated by a C_4_ concentrating mechanism. Based on the δ^13^C value in *F. palmeri*, the fixation through the C_4_ concentrating mechanism is up to 93%, which is consistent with the measured proportion of initial carbon fixation in the form of C_4_, compound (Fig. 5), suggesting at this step, the released CO_2_ in the BSCs can be largely fixed by Rubisco. Whereas, the transition between C_4_-like to C_4_ process is an evolutionarily “down-hill” process and most of optimization occurred through fine-tuning expression abundance (Figs. 1-4).

## Conclusions

Combining transcript abundance, protein sequence, morphology features, here we systematically evaluate the molecular evolutionary trajectory of C_4_ photosynthesis in the genus *Flaveria*, in particular the clade A of the genus *Flaveria*. We found a clear evolutionary coordination of different features. Our data also support evolutionary jumps during the evolution of C_4_ species, which reflect three major steps during the emergence of C_4_ metabolism, including the pre-adaptation step where GDC moved from mesophyll cell to bundle sheath cell (N3), formation of C_2_ nitrogen re-balancing pathway and concurrent formation of a C_4_ pathway (N6), and dominance of C_4_ metabolic pathway (N7). The modification at N7 shows the biggest jump during the emergence of C_4_ metabolism.

## Methods

### Data retrieval

RNA-Seq data of *Flaveria* species were downloaded from the Sequence Read Achieve (SRA) of the National Center for Biotechnology Information (NCBI) (Supplementary Methods). All accession numbers for RNA-Seq data are shown in Table S1.

CO_2_ compensation points (Γ) (except for *F. kochiana*), δ^13^C (except for *F.kochiana*), %O_2_ inhibition of P_max_ (except *F. kochiana*), and CO_2_ assimilation rates were from [23]. Γ, δ^13^and %O_2_ inhibition of *F. kochiana* were from [38]. Data for % initial C_4_ products in total fixed carbon were from [50]. Data for PWUE, PNUE, and net CO_2_ assimilation rate (*A*) versus Rubisco content were from [38]. Data for M area, M:BS ratio, vein density and number of ground tissue layers were from [14]. The values of M area, M:BS ratio and vein density were measured from figures in McKown and Dengler [14] with GetData (http://www.getdata-graph-digitizer.com). Mean values from five measurements were used. Ultrastructural data of BS cell chloroplasts were from Nakamura *et al*. [40].

### Transcriptome assembly and quantification

Transcripts of *Flaveria* species generated with Illumina sequencing were assembled using Trinity (version 2.02) [51] with default parameters (Table S1). Contigs of four *Flaveria* species from 454 sequencing data were assembled using CAP3 [52] with default parameters. In all cases, only contigs of at least 300 bp in length were saved. Transcript abundances of 31 *Flaveria* samples were analyzed by mapping Illumina short reads to assembled contigs of corresponding species and then normalized to the fragment per kilobase of transcript per million mapped reads (FPKM) using the RSEM package (version 1.2.10) [53]. Functional annotations of *Flaveria* transcripts were determined by searching for the best hit in the coding sequence (CDS) dataset of *Arabidopsis thaliana* (Arabidopsis) in TAIR 10 (http://www.arabidopsis.org) by using BLAST in protein space with E-value threshold 0.001. If multiple contigs shared the same best hit in CDS reference of Arabidopsis, then the sum FPKM of those contigs was assigned to the FPKM value of the gene in *Flaveria*. To make the FPKM comparable across different samples, we normalized the FPKM value by a scaling strategy as used by Brawand *et al*. [54]. Specifically, among the transcripts with FPKM values ranking in 20%-80% region in each sample, we identified the 1000 genes that had the most-conserved ranks among 29 leaf samples, which were then used as an internal reference, and the transcript of each sample was normalized according to the mean value of these 1000 genes in the sample. We then multiplied all the FPKM values in all samples by the mean value of 1000 genes in the 29 leaf samples (Fig. S2). Genes showing differential expression were identified by applying dexus (version 1.2.2) [55] in R, with a P value less than 0.05.

### Protein divergence, gene expression divergence and morphology divergence

Pair-wise protein divergence (dN) was calculated by applying codeml program in PAML package [56] by using F3X4 condon frequency. The input super CDS sequence was from the linked coding sequences (CDS) as used in construct phylogenetic tree of *Flaveria* genus [21], which contains 2462 genes. Gene expression divergence was calculated as Euclidean distance applying R package based on gene expression values (FPKM) of 1,2218 genes. Encoded morphology values of 30 morphology traits were from [41]. The morphology divergence was calculated as Euclidean distance of morphology values. Expression and morphology values were normalized using quantile normalization applying preprocessCore package in R. Linear regression of pair-wise correlation was inferred apply lm function in R package.

### Relative difference of each ancestral node in the phylogenetic tree

The morphological characteristics, gene expression abundance, and protein sequences at the whole transcriptomic scale were predicted using FASTML [57]. The protein alignment was from [21]. Gene expression abundance and morphological characteristics of all ancestral nodes were predicted by applying ape package of R which uses a maximal likelihood method. For all C_4_ related gene expression, protein sequences and physiological data, their values of the ancestral nodes were assigned to those of the most recent species derived from the node.

Relative difference of protein sequence at each ancestral node was inferred by comparing the sequence at this node (N) with the nearest preceding node of N (N[pre]), *e.g.*, the number of different amino acid between node2 (N2) with N1 is the number of changed amino acid at N2. The number of different amino acid changes divided by the aligned length of the protein was calculated as relative protein difference for each gene. Relative difference of gene expression and morphology were calculated as (N-N[pre])/N[pre]. In most cases, the nearest preceding node of N[i] is N[i-1], there are two exceptions: the ancestral node of N11 is N5, and N10 is N8 (Fig. 7D). One-way ANOVA analysis followed by Tukey’s Post Hoc test was used to calculate the significance of relative difference between any two ancestral nodes. *P* values were adjusted by *Benjamin-Hochberg* (BH) correction.

## List of Abbrevations

*A*: CO2 assimilation rate
AAP6: protein and amino acid permease 6
AlaAT: Alanine aminotransferase
AspAT5: aspartate aminotransferase 5
BSCs: bundle sheath cells
CET: cyclic electron transport
Cytb561: cytochrome b561
DE: differentially expressed
FPKM: fragments per kilobase of transcript per million mapped reads
GDC: glycine decarboxylase complex
GLYK: glycerate kinase
GOGAT: glutamine synthetase and glutamine oxoglutarate aminotransferase
GR: glutathione reductase
GSL1: glutamine synthetase-like 1
Lhca5: PSI light harvesting complex gene 5
MATE: multidrug and toxic compound extrusion
MCs: mesophyll cells
NADP-ME: NADP-dependent malic enzyme
NCBI: National Center for Biotechnology Information
Ndh: NADH dehydrogenase-like
NHD1: sodium: hydrogen (Na+/H+) antiporter 1
PEPC: phosphoenolpyruvate carboxylase
PIFI: post-illumina chlorophyll fluorescence increase
PNUE: instantaneous photosynthetic nitrogen use efficiency
PPCKA: PEPC protein kinase A
PPDK-RP: PPDK regulatory protein
PPDK: pyruvate, orthophosphate dikinase
PSI: photosystem I
PWUE: instantaneous photosynthetic water use efficiency
RAC: Rubisco activase
Rubisco: ribulose-1,5-bisphosphate carboxylase/oxygenase
SHM: hydroxymethyltransferase
SorDH: sorbitol dehydrogenase
SRA: Sequence Read Achieve
Γ: CO2 compensation point

## Acknowledgement

The authors thank Haiyang Hu for great suggestion, and thank Chinese Academy of Sciences and the Max Planck Society for support. This work was sponsored by Shanghai Sailing Program [17YF421900], National Science Foundation of China [31701139 to Ming-Ju Amy Lyu, 30970213 to Xin-Guang Zhu]; Bill & Melinda Gates Foundation [OPP1014417 to Xin-Guang Zhu]. We thank LetPub (www.letpub.com) for its linguistic assistance during the preparation of this manuscript. All authors declared not conflict of interest.

## Competing interests

None of the authors have any competing interests.

## Availability of data and materials

RNA-Seq used in this study is downloaded from Sequence Read Achieve (SRA) of the National Center for Biotechnology Information (NCBI), the accession number is showed in Table S1.

## Authors’ contributions

MAL, UG, YT, HC and SC conducted the analysis and wrote the paper. PW, SK, JMH, RFS, ML, GKS and XGZ designed the study and wrote the paper.

## Supplementary Information

Additional file 1: Includes supplemental methods, figures and tables

Additional file 2: The alignments of proteins

Additional file 3: 205 genes that showed differential expression and at least one amino acid change between C_3_ and C_4_ species. The table includes the gene identifier, expression fold-change and number of amino acid changes between C_4_ and C_3_ species.

Additional file 4: FPKM of all genes

## Reference

1 Pattin KA, Moore JH: Genome-wide association studies for the identification of biomarkers in metabolic diseases. Expert opinion on medical diagnostics 2010, 4(1): 39-51.

2 Manolio TA, Collins FS, Cox NJ, Goldstein DB, Hindorff LA, Hunter DJ, McCarthy MI, Ramos EM, Cardon LR, Chakravarti A et al: Finding the missing heritability of complex diseases. Nature 2009, 461(7265): 747-753.

3 Huang X, Han B: Natural variations and genome-wide association studies in crop plants. Annual review of plant biology 2014, 65: 531-551.

4 Zhu X-G, Shan L, Wang Y, Quick WP: C4 Rice - an ideal arena for systems biology research. Journal of Integrative Plant Biology 2010, 52(8): 762-770.

5 Sage RF, Christin PA, Edwards EJ: The C-4 plant lineages of planet Earth. J Exp Bot 2011, 62(9): 3155-3169.

6 Hatch MD: C4 photosynthesis: a unique blend of modified biochemistry, anatomy and ultrastructure. Biochimica et Biophysica Acta 1987, 895: 81-106.

7 Sage RF: The evolution of C4 photosynthesis. New Phytologist 2003, 161(2): 341-347.

8 Hatch MD, Slack CR: A new enzyme for the interconversion of pyruvate and phosphopyruvate and its role in the C4 dicarboxylic acid pathway of photosynthesis. The Biochemical journal 1968, 106(1): 141-146.

9 Johnson HS, Hatch MD: The C4-dicarboxylic acid pathway of photosynthesis. Identification of intermediates and products and quantitative evidence for the route of carbon flow. The Biochemical journal 1969, 114(1): 127-134.

10 Dengler N, Nelson T: Leaf structure and development in C4 plants. In.: Sage, R, F, Monson, R, K ed (s). C4 plant biology. Academic Press: San Diego, etc; 1999.

11 Hatch MD, Osmond CB: Compartmentation and transport in C4 photosynthesis. Encyclopedia of Plant Physiology 1976, 3: 144-184.

12 Slack CR, Hatch MD, Goodchild DJ: Distribution of enzymes in mesophyll and parenchyma-sheath chloroplasts of maize leaves in relation to the C4-dicarboxylic acid pathway of photosynthesis. The Biochemical journal 1969, 114(3): 489-498.

13 Sage RF, Sage TL, Kocacinar F: Photorespiration and the evolution of C4 photosynthesis. Annual review of plant biology 2012, 63: 19-47.

14 McKown AD, Dengler NG: Key innovations in the evolution of Kranz anatomy and C4 vein pattern in Flaveria (Asteraceae). American journal of botany 2007, 94(3): 382-399.

15 Brown NJ, Parsley K, Hibberd JM: The future of C4 research--maize, Flaveria or Cleome? Trends in plant science 2005, 10(5): 215-221.

16 Gowik U, Brautigam A, Weber KL, Weber AP, Westhoff P: Evolution of C4 photosynthesis in the genus Flaveria: how many and which genes does it take to make C4? Plant Cell 2011, 23(6): 2087-2105.

17 Mallman J, Heckmann D, Brautigam A, Lercher MJ, Webb APM, Westhoff P, Gowik U: The role of photorespiration during the evolution of C4 photosynthesis in the genus Flaveria. eLife 2014, 3:e02478.

18 Engelmann S, Blasing OE, Gowik U, Svensson P, Westhoff P: Molecular evolution of C4 phosphoenolpyruvate carboxylase in the genus Flaveria--a gradual increase from C3 to C4 characteristics. Planta 2003, 217(5): 717-725.

19 Westhoff P, Gowik U: Evolution of c4 phosphoenolpyruvate carboxylase. Genes and proteins: a case study with the genus Flaveria. Annals of botany 2004, 93(1): 13-23.

20 Engelmann S, Wiludda C, Burscheidt J, Gowik U, Schlue U, Koczor M, Streubel M, Cossu R, Bauwe H, Westhoff P: The gene for the P-subunit of glycine decarboxylase from the C4 species Flaveria trinervia: analysis of transcriptional control in transgenic Flaveria bidentis (C4) and Arabidopsis (C3). Plant physiology 2008, 146(4): 1773-1785.

21 Lyu MJ, Gowik U, Kelly S, Covshoff S, Mallmann J, Westhoff P, Hibberd JM, Stata M, Sage RF, Lu H et al: RNA-Seq based phylogeny recapitulates previous phylogeny of the genus Flaveria (Asteraceae) with some modifications. BMC evolutionary biology 2015, 15(1): 116.

22 Edwards GE, Ku MS: Biochemistry of C3-C4 intermediates. In: Hatch MD, Boardman NK, editors. The biochemistry of plants New York: Academic Press 1987: 275-325.

23 Ku MS, Wu J, Dai Z, Scott RA, Chu C, Edwards GE: Photosynthetic and photorespiratory characteristics of flaveria species. Plant physiology 1991, 96(2): 518-528.

24 Matsuoka M, Tada Y, Fujimura T, Kano-Murakami Y: Tissue-specific light-regulated expression directed by the promoter of a C4 gene, maize pyruvate, orthophosphate dikinase, in a C3 plant, rice. Proceedings of the National Academy of Sciences of the United States of America 1993, 90(20): 9586-9590.

25 Schaffner AR, Sheen J: Maize C4 photosynthesis involves differential regulation of phosphoenolpyruvate carboxylase genes. The Plant journal: for cell and molecular biology 1992, 2(2): 221-232.

26 Li Y, Xu J, Haq NU, Zhang H, Zhu XG: Was low CO2 a driving force of C4 evolution: Arabidopsis responses to long-term low CO2 stress. J Exp Bot 2014, 65(13): 3657-3667.

27 Aubry S, Kelly S, Kumpers BM, Smith-Unna RD, Hibberd JM: Deep evolutionary comparison of gene expression identifies parallel recruitment of trans-factors in two independent origins of C4 photosynthesis. PLoS genetics 2014, 10(6):e1004365.

28 John CR, Smith-Unna RD, Woodfield H, Covshoff S, Hibberd JM: Evolutionary convergence of cell-specific gene expression in independent lineages of C4 grasses. Plant physiology 2014, 165(1): 62-75.

29 Chang YM, Liu WY, Shih AC, Shen MN, Lu CH, Lu MY, Yang HW, Wang TY, Chen SC, Chen SM et al: Characterizing regulatory and functional differentiation between maize mesophyll and bundle sheath cells by transcriptomic analysis. Plant physiology 2012, 160(1): 165-177.

30 Tausta SL, Li P, Si Y, Gandotra N, Liu P, Sun Q, Brutnell TP, Nelson T: Developmental dynamics of Kranz cell transcriptional specificity in maize leaf reveals early onset of C4-related processes. Journal of experimental botany 2014, 65(13): 3543-3555.

31 Ferro M, Salvi D, Brugiere S, Miras S, Kowalski S, Louwagie M, Garin J, Joyard J, Rolland N: Proteomics of the chloroplast envelope membranes from Arabidopsis thaliana. Molecular & Cellular Proteomics 2003, 2(5): 325-345.

32 Das S, Pathak R, Choudhury D, Raghuram N: Genomewide computational analysis of nitrate response elements in rice and Arabidopsis. Mol Genet Genomics 2007, 278(5): 519-525.

33 Rentsch D, Hirner B, Schmelzer E, Frommer WB: Salt stress-induced proline transporters and salt stress-repressed broad specificity amino acid permeases identified by suppression of a yeast amino acid permease-targeting mutant. The Plant Cell Online 1996, 8(8): 1437-1446.

34 Okumoto S, Schmidt R, Tegeder M, Fischer WN, Rentsch D, Frommer WB, Koch W: High Affinity Amino Acid Transporters Specifically Expressed in Xylem Parenchyma and Developing Seeds of Arabidopsis. Journal of Biological Chemistry 2002, 277(47): 45338-45346.

35 Hunt E, Gattolin S, Newbury HJ, Bale JS, Tseng H-M, Barrett DA, Pritchard J: A mutation in amino acid permease AAP6 reduces the amino acid content of the Arabidopsis sieve elements but leaves aphid herbivores unaffected. J Exp Bot 2010, 61(1): 55-64.

36 Allen JF, Pfannschmidt T: Balancing the two photosystems: photosynthetic electron transfer governs transcription of reaction centre genes in chloroplasts. Philos Trans R Soc Lond Ser B-Biol Sci 2000, 355(1402):1351-1357.

37 Foyer C, Noctor G: Redox regulation in photosynthetic organisms: signaling, acclimation, and practical implications. Antioxid Redox Signal 2009, 11(4): 861-905.

38 Vogan PJ, Sage RF: Water-use efficiency and nitrogen-use efficiency of C(3)-C(4) intermediate species of Flaveria Juss. (Asteraceae). Plant, cell & environment 2011, 34(9): 1415-1430.

39 Rumpho ME, Ku MS, Cheng SH, Edwards GE: Photosynthetic Characteristics of C(3)-C(4) Intermediate Flaveria Species : III. Reduction of Photorespiration by a Limited C(4) Pathway of Photosynthesis in Flaveria ramosissima. Plant physiology 1984, 75(4): 993-996.

40 Nakamura N, Iwano M, Havaux M, Yokota A, Munekage YN: Promotion of cyclic electron transport around photosystem I during the evolution of NADP-malic enzyme-type C4 photosynthesis in the genus Flaveria. The New phytologist 2013, 199(3): 832-842.

41 McKown AD, Moncalvo J-M, Dengler NG: Phylogeny of Flaveria (Asteraceae) and inference of C4 photosynthesis evolution. American Journal of Botany 2005, 92(11): 1911-1928.

42 Sage RF, Zhu X-G: Exploiting the engine of C4 photosynthesis. Journal of experimental botany 2011, 62(9): 2989-3000.

43 Wang Y, Long SP, Zhu XG: Elements required for an efficient NADP-malic enzyme type C4 photosynthesis. Plant physiology 2014, 164(4): 2231-2246.

44 Wedding RT, Black MK, Meyer CR: Inhibition of phosphoenolpyruvate carboxylase by malate. Plant physiology 1990, 92(2): 456-461.

45 Wang S, Tholen D, Zhu XG: C4 photosynthesis in C3 rice: a theoretical analysis of biochemical and anatomical factors. Plant, cell & environment 2017, 40(1): 80-94.

46 Hunt BG, Ometto L, Keller L, Goodisman MAD: Evolution at Two Levels in Fire Ants: The Relationship between Patterns of Gene Expression and Protein Sequence Evolution. Mol Biol Evol 2013, 30(2): 263-271.

47 Warnefors M, Kaessmann H: Evolution of the Correlation between Expression Divergence and Protein Divergence in Mammals. Genome Biol Evol 2013, 5(7): 1324-1335.

48 Morgan CL, Turner SR, Rawsthorne S: Coordination of the Cell-Specific Distribution of the 4 Subunits of Glycine Decarboxylase and of Serine Hydroxymethyltransferase in Leaves of C3-C4 Intermediate Species from Different Genera. Planta 1993, 190(4): 468-473.

49 Sage TL, Busch FA, Johnson DC, Friesen PC, Stinson CR, Stata M, Sultmanis S, Rahman BA, Rawsthorne S, Sage RF: Initial events during the evolution of C4 photosynthesis in C3 species of Flaveria. Plant physiology 2013, 163(3): 1266-1276.

50 Moore Bd, Ku M, S. B., Edwards G, E.: C4 photosynthesis and light-dependent accumulation of inorganic carbon in leaves of C3-C4 and C4 Flaveria species. Australian Journal of Plant Physiology 1987, 14: 658-668.

51 Grabherr MG, Haas BJ, Yassour M, Levin JZ, Thompson DA, Amit I, Adiconis X, Fan L, Raychowdhury R, Zeng Q et al: Full-length transcriptome assembly from RNA-Seq data without a reference genome. Nature biotechnology 2011, 29(7): 644-652.

52 Huang X, Madan A: CAP3: A DNA sequence assembly program. Genome research 1999, 9(9): 868-877.

53 Li B, Dewey CN: RSEM: accurate transcript quantification from RNA-Seq data with or without a reference genome. BMC bioinformatics 2011, 12:323.

54 Brawand D, Soumillon M, Necsulea A, Julien P, Csardi G, Harrigan P, Weier M, Liechti A, Aximu-Petri A, Kircher M et al: The evolution of gene expression levels in mammalian organs. Nature 2011, 478(7369): 343-348.

55 Klambauer G, Unterthiner T, Hochreiter S: DEXUS: identifying differential expression in RNA-Seq studies with unknown conditions. Nucleic acids research 2013, 41(21):e198.

56 Yang Z: PAML: a program package for phylogenetic analysis by maximum likelihood. Computer applications in the biosciences : CABIOS 1997, 13(5): 555-556.

57 Ashkenazy H, Penn O, Doron-Faigenboim A, Cohen O, Cannarozzi G, Zomer O, Pupko T: FastML: a web server for probabilistic reconstruction of ancestral sequences. Nucleic acids research 2012, 40(Web Server issue):W580-584.

